# Has the time come for preprints in biology?

**DOI:** 10.1101/045799

**Authors:** Needhi Bhalla

## Abstract

Preprints, non-peer reviewed drafts of manuscripts available on the internet, have been used in conjunction with peer review and publication in journals in the physical sciences for almost 25 years. Recently, more scientists have been discussing whether preprints can play a similar role in biological and biomedical research. Here, I discuss my excitement and concerns about the role that preprints can play in disseminating research findings in the life sciences.

## Introduction

On February 16^th^ and 17^th^, 2016, a small group of biologists, publishers, and funders met at a workshop on Accelerating Science and Publication in Biology (ASAPbio) at the Howard Hughes Medical Institute in Chevy Chase, Maryland. The objective of the workshop was to discuss the future of preprints in biological and biomedical publishing. Organized by Daniel Colón Ramos, Jessica Polka, Ron Vale, and Harold Varmus, this workshop attempted to identify barriers that have prevented the use of preprints in biomedical research and to determine how those barriers might be overcome to promote greater use of preprints. Similar to how preprints work in other fields, the organizers of the meeting were firmly committed to the idea that posting of preprints should be followed by publication in peer-reviewed journals. I participated in this discussion and came away very hopeful about the use of preprints. Nonetheless, I also had some concerns about preprints and how they would be viewed by members of the research community.

## What are preprints?

Preprints are non-peer-reviewed drafts of research papers posted on the internet. Perhaps the most well-known repository for preprints is arXiv, where a proportion of physicists, mathematicians and astronomers have been posting their papers for almost 25 years. 64% of preprints posted on arXiv are subsequently published in peer-reviewed journals (Lariviere *et al*., 2014). This is likely an underestimate given the delay between posting on arXiv and publication. Indeed, if similar analysis is limited to articles posted on arXiv between 1995 and 2006, 73% of preprints are published in peer-reviewed journals (Lariviere *et al*., 2014). Thus, posting a preprint on arXiv does not replace peer-review or publication in journals but exists alongside them. Because of this co-existence, most journals that publish physics, mathematics or astronomy papers have editorial review policies indicating that they are receptive to evaluating papers that have already been posted on arXiv.

The use of arXiv varies tremendously across disciplines and is by no means the norm. Only 20% of published papers in the general field of physics also appear on arXiv (Lariviere *et al*., 2014). This reflects the fact that the use of arXiv is standard in some subfields and atypical in others. For example, posting on arXiv accounts for 60-70% of published articles in astronomy, astrophysics and nuclear and particle physics (Lariviere *et al*., 2014). However, authors in other fields, such as solid state physics, post preprints on arXiv as well as submit them to journals about 30% of the time (Lariviere *et al*., 2014). Therefore, the decision to post a preprint on arXiv is likely a deliberate choice and not a foregone conclusion.

For the biomedical sciences, there are a variety of models for preprints, including arXiv, biorXiv, PeerJ and F1000Research. Posting preprints in biology is rapidly accelerating and appears to be more common in certain fields, such as evolutionary biology, bioinformatics and genomics (Inglis and Sever, 2016). It was in this context that ASAPbio was organized to address what role preprints could play in publishing biomedical research. The focus of ASAPbio was to discuss how preprints could be used in conjunction with peer review and journal publication, similar to their use in physics, mathematics and astronomy. The idea proposed was that preprints would be posted simultaneously or soon after a manuscript was submitted to a journal. The hope was that this approach could help deal with one very specific current concern about scientific publishing: the delay, often substantial, that exists between having a completed manuscript that documents a scientific study, and its subsequent publication and dissemination by journals.

## What are the advantages of preprints?

The greatest advantage that preprints afford is the ability to disseminate one’s research rapidly when the scientists involved have decided that the study is largely complete. The ability to let other scientists know of recent developments in your lab so that they can rapidly build and expand upon those developments will promote more rapid progress in a field than the fits and starts that often accompany the peer review process for journals. In addition, posting a preprint could establish priority of discovery, identifying your work as amongst the first to demonstrate a research finding.

Practically, preprints can demonstrate productivity and scholarly contributions to a field while a manuscript is being peer-reviewed and vetted for publication in a journal. Having a completed manuscript that is readily accessible on the internet, even when under review at a journal, is far more tangible than the tenuous assertion on a CV that it is in preparation or has been submitted to a journal. Thus, preprints could demonstrate completion of studies without the delays associated with publication. Trainees could use preprints to demonstrate productivity as they prepare for career transitions, while faculty members could use preprints in assembling promotion dossiers or grant applications. Members of tenure and promotion committees, grant reviewers and employers would be able to assess the manuscript and make professional judgments that are not held hostage to the manuscript review and publication process that can stretch for months, if not years.

Preprints provide additional opportunities. One’s work becomes more widely disseminated earlier through preprints. If a manuscript is ultimately published in a pay-walled journal, individuals without journal subscriptions can at least access the manuscript in its preprint form. Other researchers interested in the study can comment and review the manuscript in its preprint form while it is also being reviewed at a journal, potentially generating a stronger, more rigorous study because it has been evaluated by more than the 2-4 scientists that participated in a journal’s review process.

## What are the risks associated with preprints?

One of the greatest challenges associated with preprints is the uncertainty about how they will be perceived and evaluated by other scientists. Will other scientists in my field acknowledge preprints as evidence of productivity, scholarly contributions to a field and priority of discovery? Will they acknowledge evidence presented in a preprint as the first demonstration of a research finding, for example, by citing the preprint, even if eventual publication in a peer reviewed journal is significantly delayed? Is there a danger of getting “scooped” if I post a preprint? Will preprints be evaluated and respected as a first step towards publication in a peer-reviewed journal? Which peer-reviewed journals will reject a manuscript if it has previously appeared as a preprint?

All of these concerns are exacerbated by the current funding climate. As scientists, our job is to perform research, disseminate our discoveries, and mentor the next generation of scientists. All of these require funding. Many investigators are perpetually worried about how to keep their labs funded. The uncertainty about how preprints will be viewed by funding agencies, tenure committees, journal editors and other scientists in the field may present too great a risk. This is likely to be especially true for new investigators. These concerns can also infect our trainees, some of whom would like to continue in academic science. They may be skeptical of demands for change by faculty members who have already prospered in the current system.

I have discussed preprints with other scientists who did not attend the ASAPbio workshop. Most have not even considered posting preprints. Among those who have, some embrace the concept and leave me convinced that risks are exaggerated. Others have heard about preprints and perceive them negatively, sometimes lumping them with low-impact papers published in predatory journals. Often, these scientists appreciate the gatekeeper roles that well-established journals play and worry about the quality of science that will get posted as preprints but may never be subsequently published in a journal. Studies on preprints that have been posted on arXiv and subsequently published in journals argue against the claim that the availability of posting preprints will result in an explosion of low-impact studies (Davis and Fromerth, 2007; Gentil-Beccot *et al*., 2009; Lariviere *et al*., 2014). Given the potential implications for human health, some might argue that comparing the standards for disseminating research in the physical sciences with those of biomedical research is inaccurate. Unfortunately, we know of numerous examples of peer-reviewed, published research that were ultimately found to be incorrect or fabricated, demonstrating that peer-review in and of itself is not a complete safeguard against low quality publications. Despite the fact that I don’t agree with these negative perceptions of preprints, I have to consider that these scientists may review my grants, my papers and my promotions and factor that substantial risk against the advantages that I easily recognize.

Coupled with this uncertainty is the absence of obvious structural support for preprints. Granting agencies, university promotion and tenure committees and some publishers do not have clear and transparent policies regarding how preprints should evaluated. For example, some journals accept manuscripts that have been posted as preprints, while others that are well regarded in my field have policies either completely incompatible or potentially compatible with the posting of preprints (for journal compatibility with preprints, see: https://en.wikipedia.org/wiki/List_of_academic_journals_by_preprint_policy). This lack of structural support and clear policy further reinforces the idea that scientists cannot predict how other scientists will evaluate their preprints. For example, if a funding agency does not have a policy on how preprints will be evaluated during grant review and an individual reviewing a grant has a negative opinion about preprints, will that affect the review of a grant that includes references to preprints? If there is a well-defined NIH or NSF policy explaining how to assess preprints, the Scientific Review Officer or other members of the study section have a firm foundation from which they can guide the discussion to prevent bias from negatively affecting the review of grants.

## Performing a personal calculus: Under what circumstances would I feel comfortable using preprints to disseminate scientific research?

After attending ASAPbio and thinking deeply about preprints, I am absolutely intrigued by and optimistic about the opportunities provided by preprints. However, I am also concerned enough about potential disadvantages to assess the risk of posting a preprint with every manuscript that my lab is planning to submit to a journal. In addition to the concerns addressed above, I would also consider whether community access to the research findings in a particular manuscript is time-sensitive or relevant to immediate public health initiatives. What journal(s) would I like to submit my paper to and what is their policy towards reviewing preprints? Do my trainees feel comfortable submitting a preprint? Am I, or the members of my lab, concerned about being scooped and would we therefore prefer to establish priority with a journal article instead of a preprint?

These calculations might be different for each manuscript. Considering the analysis of preprint posting by members of the physics, astronomy and mathematics communities

(Lariviere *et al*., 2014), this personal calculus would appear to be the norm even amongst those who use arXiv.

In addition, it has become increasingly clear after the ASAPbio meeting that we need to discuss with other scientists what preprints mean for biological and biomedical research. If a major concern about preprints is that we will lose the valuable role peer-review plays in improving manuscripts, we should remind people that posting a preprint is likely only the first step to ultimately publishing a paper in a journal after peer-review. If a major impediment to posting preprints is the uncertainty about how others perceive preprints, then initiating conversations about preprints could assuage any concerns. These are conversations that should occur with members of one’s department, the administration at one’s university, colleagues in one’s field and granting agencies that fund the research in one’s lab. Moreover, these discussions could help influence policy so that structural support for preprints soon follows, providing institutional guidelines that can help further develop the support of individual scientists. After all, we are the scientists that review grants in study sections, evaluate and write letters for promotion dossiers and make up faculty search committees. By instigating these conversations, we can begin to build consensus for how we will assess preprints so we can take advantage of the important opportunities they present.

## Acknowledgements

I would like to thank Darren Boehning for bringing “arXiv E-prints and the journal of record: An analysis of roles and relationships” to my attention and Doug Kellogg for critical reading of this editorial.

